# Age-associated modulation of TREM1/2- expressing macrophages promotes melanoma progression and metastasis

**DOI:** 10.1101/2024.11.20.624563

**Authors:** Marzia Scortegagna, Rabi Murad, Parinaz Bina, Yongmei Feng, Rebecca Porritt, Alexey Terskikh, Xiao Tian, Peter D. Adams, Kristiina Vuori, Ze’ev A. Ronai

## Abstract

Aging is a known risk factor for melanoma, yet mechanisms underlying melanoma progression and metastasis in older populations remain largely unexplored. Among the current knowledge gaps is how aging alters phenotypes of cells in the melanoma microenvironment. Here we demonstrate that age enriches the immunosuppressor tumor microenvironment, which is linked to phenotypes associated with melanoma metastasis. Among cellular populations enriched by aging were macrophages with a tolerogenic phenotype expressing TREM2 and dysfunctional CD8-positive cells with an exhausted phenotype, while macrophages with profibrotic phenotype expressing TREM1 were depleted. Notably, TREM1 inhibition decreased melanoma growth in young but not old mice, whereas TREM2 inhibition prevented lung metastasis in aged mice. These data identify novel targets associated with melanoma metastasis and may guide aged-dependent immunotherapies.

## Introduction

Aging is among the main risk factors for cancer development and progression^1,2^. Accordingly, in melanoma, as in other tumor types, the incidence of invasive tumors increases with age^3,4,5^. Yet, despite the clear link between aging and cancer, most preclinical studies of melanoma have not considered the role played by aging in disease progression. The average age of melanoma mouse models is 6-8 weeks (equivalent to 15 to 20-year-old humans), highlighting the need for models that better recapitulate biological changes impacting cancer progression in older individuals. Increased incidence of driver mutations seen in older patients with melanoma is associated with genomic instability in cancer cells^6,7^. Aging is also known to be associated with low-grade chronic inflammation, which promotes cancer development^8,9^. By contrast, how the aged tumor environment contributes to melanoma progression and invasiveness is less understood. Melanoma differs in older age groups, with progressively higher rates of metastasis and death. These clinical data support the need to understand the increased propensity to metastasis and diminished response to immunotherapy in the aged population.

Aged fibroblasts were recently shown to be capable of promoting melanoma metastasis and therapy resistance^10^. In considering the potential importance of immune cell components to melanoma metastasis, we focused on macrophages, which are among the most abundant immune cells infiltrating melanoma tumors and are implicated in tumor progression and metastasis due to their effect on cancer cell proliferation, angiogenesis, and anti-tumor immunity^11,12,13^. Macrophages are a heterogeneous population of innate immune cells with several functions, including phagocytosis of pathogens, clearance of dead cells, and cytokine production^14^. While initially classified as pro-(M1) or anti-(M2) inflammatory^15^, single-cell RNA-seq studies have revealed that tumor-associated macrophages (TAMs) express a mix of M1- and M2-like genes and have identified tissue- and stimuli-specific macrophage subsets^16,17,18,19,20^. Decreases in phagocytosis and chemotaxis, both associated with impaired resolution of inflammation, have been seen in macrophages of aged individuals^21,22^. Yet the role of macrophages in melanoma progression in the elderly population remains largely unknown.

In this study, we examined age-associated changes in immune cells infiltrating melanoma tumors and showed that changes seen in macrophages contribute to age-dependent melanoma progression and metastasis. These observations provide important insight into how the tumor microenvironment may change with aging and the importance of macrophages in melanoma metastasis, which is commonly seen in the aged population.

## Results

### Aging promotes lung metastasis and alters the composition of immune cells in melanoma tumors’

To evaluate the role of aging in melanoma progression and invasiveness, we inoculated 4-, 12- and 20-month (M) old mice, representing respective young, middle-aged, and aged mice, with YUMM1.7 melanoma cells^23,24^. We observed decreased tumor growth in 20M old mice, relative to the 12M or 4M month-old mice (Fig. 1A and B). Notably, we observed a significant increase in lung metastasis in 12M and 20M mice, relative to 4M mice (Fig. 1C, and extended data Fig. 1A, 1B). These results are consistent with previous reports that melanoma becomes more invasive with age^10,2^ and support the use of aged mice as a model to define mechanisms underlying melanoma metastasis.

**Figure 1.**
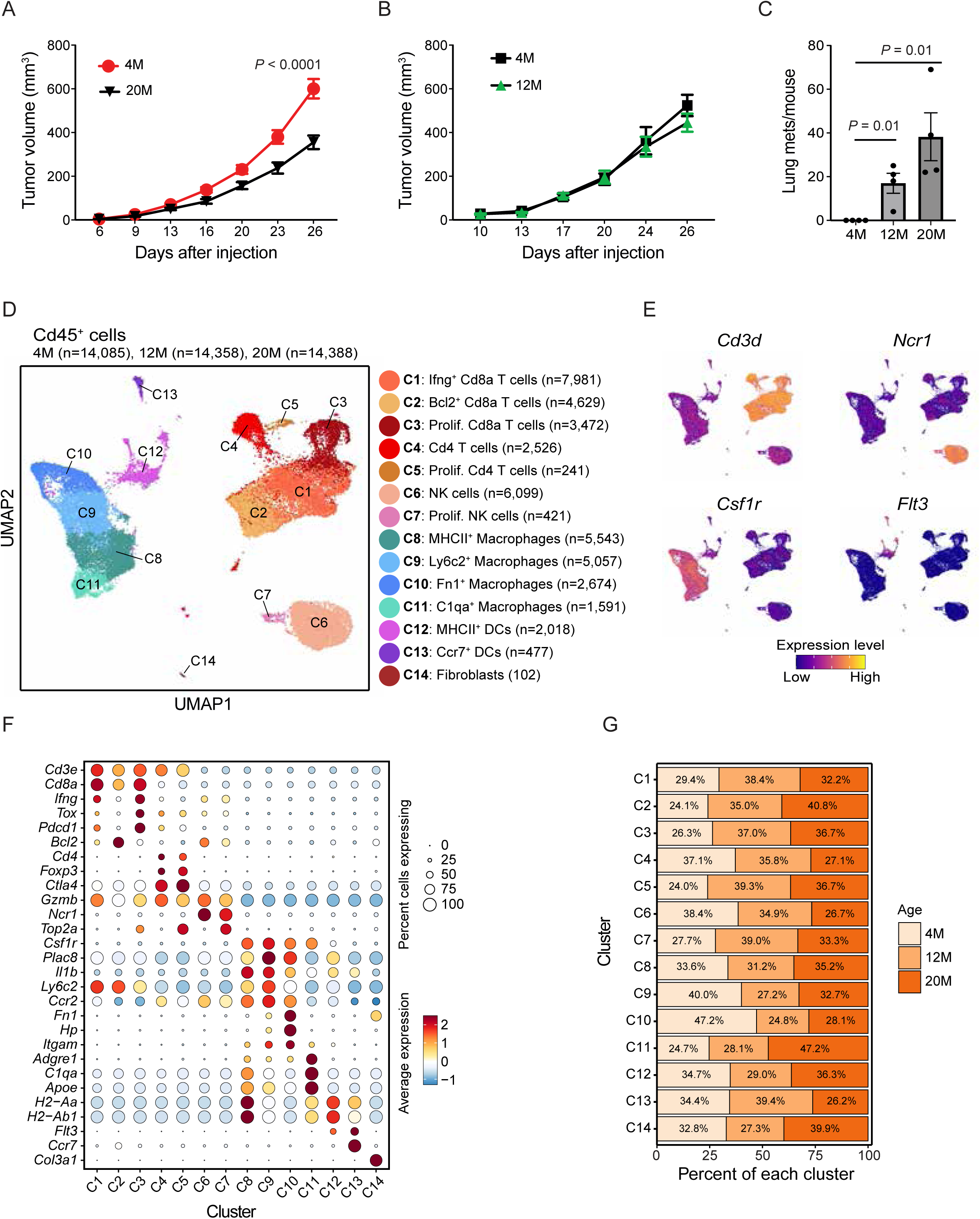
Aging promotes lung metastasis and alters composition of immune cells in melanoma tumors. (**A-B**) YUMM1.7 melanoma cells (500,000) were injected s.c. into the flank of either 4 month- (4M) and 20 month- (20M) (A) or 4 month- (4M) and 12 month- (B) old mice and tumor growth (volume) was measured over time. n = 10 for 4M, n = 8 for 20M in panel A; n = 6 for 4M, n = 5 for 12M in panel B. (**C**) YUMM1.7 melanoma cells (500,000) were injected s.c. into the flank of 4M, 12M and 20M old mice and after 22 days lungs were collected and processed to obtain 5 serial sections per lung, which were analyzed by IHC of S100, a marker of melanoma cells, to quantify S100-positive colonies (more than 10 S100^+^ cells per lesion). N = 4 for each group. (**D**) CD45^+^ cells were sorted by flow cytometry from melanoma tumors of 4M, 12M and 20M mice and single cell RNA-seq was performed. Integrated UMAP plot of CD45^+^ cells from melanoma tumors (7 tumors/ group) collected 14 days after inoculation of YUMM1.7 cells showing different immune cell clusters. Immune cell types identified based on expression of specific cell markers are labeled. (**E**) UMAP plots showing marker genes expression used to identify immune cell populations. (**F**) Dot plots showing scaled expression of marker genes used to identify specific immune cell populations. (**G**) Bar graph showing the percentages of various immune cells for each cluster from 4M, 12M and 20M old mice. Data in panels A, B and C are presented as means ± SEM. Data in panel A were analyzed by two-way ANOVA. Data in panel C were analyzed by unpaired *t*-test.

To identify age-dependent changes in tumor-infiltrating immune cell populations, we performed single-cell RNA-seq (scRNA-seq) analysis of CD45^+^ cells from YUMM1.7 tumors grown in 4M, 12M, and 20M mice. We performed integrated clustering of the three single cell datasets using Seurat. Fourteen immune cell clusters harboring distinct gene expression patterns were identified in this analysis of CD45^+^ cells of tumors isolated from 4M (n=14,085 cells), 12M (n=14,358 cells), and 20M (n=14,388 cells) mice (Fig. 1D-F; Table S1). Those included Ncr1^+^ NK cells (clusters C6 and C7), Cd3e^+^ T cells (clusters C1-C5), Csfr1^+^ myeloid cells (clusters C8-C11) and Flt3^+^ dendritic cells (DCs; clusters C12, C13). T-cell populations were distributed into effector CD8^+^ T cells that expressed high interferon γ levels (cluster C1), exhausted and proliferative CD8^+^ T cells expressing the exhaustion marker Tox and proliferative marker Top2a (cluster C3), CD4^+^ T cells expressing the Treg marker Foxp3; cluster C4) and proliferating CD4^+^ cells (cluster C5). Within the myeloid population of Csf1r^+^ cells, cluster C8 exhibited the highest MHCII expression, whereas clusters C9 and C10 were enriched in the monocytic markers Ccr2 and Plac8. Of those, cluster C9 exhibited a high level of inflammatory markers (Il1b), while cluster C10 showed a high expression of pro-fibrotic markers (Fn1 and Hp). Cluster 11 showed the highest expression of markers of mature and resident-like macrophages (C1qa and Adgre1). Of note, aging markedly changed the frequencies of cells in macrophage clusters C10 and C11 (Fig. 2G). Specifically, we observed an increase in mature macrophages (cluster C11; 47.2% vs. 24.7%) and a decrease in pro-fibrotic macrophages (cluster C10; 28.1% vs. 47.2%; Fig. 2G) in aged (20M) compared to young (4M) mice. These data suggest that phenotypic changes seen in macrophages isolated from aged mice may be linked with an increased propensity of melanoma to metastasize.

**Figure 2.**
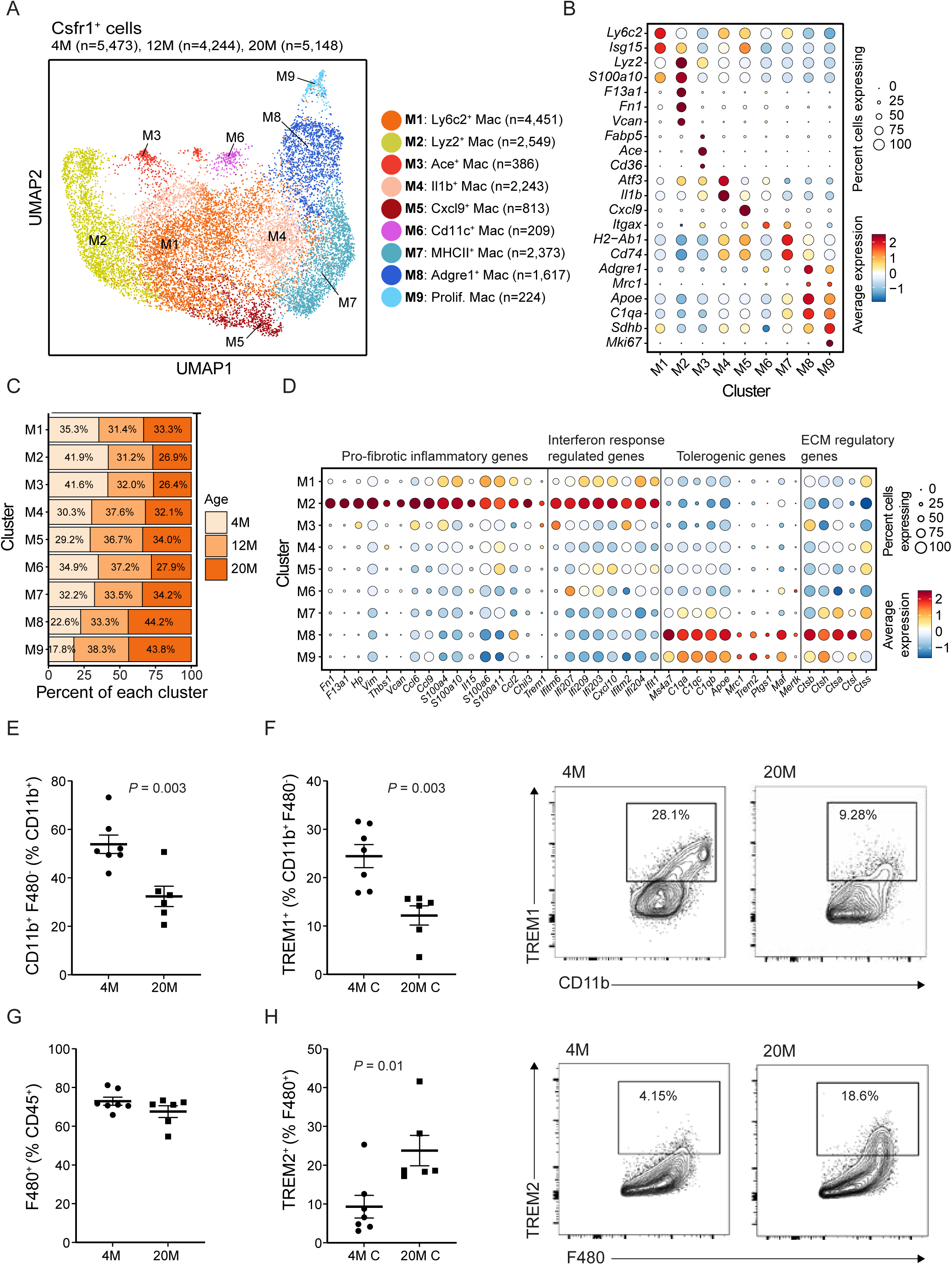
Aging is conducive to the establishment of tolerogenic macrophages in melanoma. Csfr1^+^ cells (clusters C8-C11 from global UMAP in Figure 1D) from scRNA-seq analysis of CD45^+^ cells were re-clustered to obtain higher resolution clustering of myeloid cells. (**A**) Integrated UMAP plot of myeloid cells from 4M (n= 5,473 cells), 12M (n= 4,244 cells) and 20M (n= 5,148 cells) tumors showing 9 sub-clusters. (**B**) Dot plots showing scaled expression of marker genes used to identify myeloid cell sub-clusters. (**C**) Bar graph showing the percentages of various myeloid cell sub-clusters of for each cluster from 4M, 12M and 20M old mice. (**D**) Dot plots showing scaled expression of indicated genes used to identify specific functions. (**E**) Frequencies of CD11b^+^ F480^low^ among CD11b^+^ cells from CD45^+^ cells of tumors from 4M and 20M old mice. (**F**) Frequencies of TREM1^+^ among CD11b^+^ F480^low^ cells and representative flow cytometry plots showing TREM1^+^ expression by a gated subpopulation (CD11b^+^ cells) of CD45.2^+^ cells from 4M and 20M old mice. (**G**) Frequencies of F480^+^ among CD45^+^ cells in tumors from 4M and 20M old mice. (**H**) Frequencies of TREM2^+^ among F480^+^ cells and representative flow cytometry plots showing TREM2^+^ expression by a gated subpopulation (F480^+^ cells) of CD45.2^+^ cells from 4M and 20M old mice. n = 7 for 4M and n = 6 for 20M for panels E-H. Data in panels D-H were analyzed by unpaired *t*-test.

#### Aging favors the establishment of tolerogenic macrophages in melanoma

TAMs play a key role in angiogenesis, extracellular remodeling, and immunosuppression, phenotypes conducive to increased melanoma invasiveness and metastasis^11,12, 13^. To further assess changes in age-dependent TAM signaling and assess their potential role in melanoma progression and invasiveness we examined our scRNA-seq data and re-clustered Csf1r-positive macrophages, resulting in 9 clusters (Figs. 2A and 2B; extended data Table S2). Clusters M1, M2, M4, and M5 exhibited high expression of the monocyte marker Ly6c2, coinciding with high expression of interferon response-regulated genes (Isg15) in cluster M1, profibrotic genes (Fn1, F13a1, and Vcan) in cluster M2, genes encoding inflammatory cytokines (Il1b) in cluster M4 and chemokines (Cxcl9) in cluster M5. In contrast, cluster M3 exhibited the highest expression of genes associated with lipid metabolism (Cd36, fabp5, and Ace). Cluster M7 showed the highest expression of genes associated with antigen presentation (MHCII), while cluster M6 exhibited high expression of Itgax. Macrophage clusters expressing markers of mature (Adgre1) and pro-tumorigenic (Apoe, C1qa, and Mre1) markers but not of monocyte markers (Ccr2) were identified in clusters M8 and M9. (Fig. 2B and extended data Fig. 2A). Cluster M9 contained proliferating Ki67-positive macrophages.

The most profound age-related changes in macrophage clusters were the depletion of clusters M2 and M3 and the enrichment of clusters M8 and M9 (Fig. 2C). Further analysis of markers specific to M2 revealed high expression of pro-fibrotic and inflammatory genes (such as Trem1, S100a10 and Fn1) and Interferon response regulated genes (including Ifitm6 and Ifi207; Fig. 2D), likely reflecting exposure to interferon signaling. Cluster M3, which was depleted with aging, was marked by high expression of genes involved in lipid metabolism, which were implicated in pro-tumorigenic effects of tumor-associated macrophages (extended data Fig. 2B). The notable increase in M8 and M9 clusters seen in 12M and 20M relative to 4M mice coincided with elevated expression of genes implicated in immune suppression (Apoe, Trem2, and Maf), cancer cell invasiveness (cathepsins and genes implicated in extracellular matrix remodeling; Fig. 2D), and encoding specific chemokines (Ccl4 and Ccl12; extended data Fig. 2C) in these clusters.

Among the most differentially expressed genes in age-associated cluster M8 were genes encoding the fat binding protein Apoe and the transcription factor Maf, both significantly upregulated in 20M relative to 4M mice (extended data Fig. 2D; extended data Tables S3-S11). These genes are expressed in immunosuppressive macrophages and implicated in immunosuppressive phenotypes, concomitant with increases in expression of Ctsl, a protease that functions in tumor invasiveness and metastasis^25^ (extended data Fig. 2D; extended data Tables 3-11). In contrast, the genes most upregulated in young-associated cluster M2 (from tumors in 4M relative to 20M mice) included interferon response-regulated genes (Isg20, Irf7, and Ifi203: extended data Fig. 2E), which were implicated in anti-tumor activity. Of interest, several macrophages’ clusters (including cluster M2) derived from 4M mice also expressed induction of genes functioning in cancer progression and invasiveness, such as Plaur and Spp1 (extended data Fig. 2E; extended data Tables 3-11) when compared to 20M macrophages, suggesting the existence of a heterogeneous macrophage subpopulation that expresses both immunostimulatory and immunosuppressive genes at this young age.

To confirm conclusions based on scRNA-seq analyses we monitored changes in immune cell infiltration of tumors using FACS analysis. While the frequencies of CD45^+^ cells among all cells and of CD11b^+^ cells among CD45^+^ cells were comparable in tumors from 4M and 20M mice (extended data Figs. 2F, 2G), we observed a significant increase of CD11b^+^ and F480^low^-expressing cells in CD11b-positive macrophages from 4M mice (Fig. 2E), suggesting that young mice contain less tumor infiltrated mature macrophages expressing F480 compared with the aged population. Moreover, CD11b^+^ F480^low^ immune cells in tumors from 20M mice showed a significant decrease of TREM1 expressing cells, relative to 4M mice (Fig. 2F). While we observed no age-associated changes in F480^+^ macrophages among CD45^+^ cells, the frequency of TREM2^+^ cells relative to F480^+^ macrophages significantly increased with age (comparing 20M with 4M old mice; Figs. 2G and 2H). Of note, the frequency of CD36^+^ cells among CD11b^+^ cells was decreased in 20M relative to 4M mice (extended data Fig. 2H). Overall, these findings support conclusions drawn from our scRNA-seq analysis indicating an overall decrease in TREM1-positive macrophages and an increase in TREM2-positive, mature macrophages in aged relative to young mice. Additional comparisons between 4M and 12M old mice confirmed these observations and indicated a decrease in TREM1-expressing cells among CD11b^+^ cells infiltrating tumors plus an increase of MRC1-expressing cells in F480^+^ macrophages (extended data Fig. 2I). Of note, MRC1 expression was specifically identified in age associated macrophage clusters M8 and M9 from scRNA-seq analysis (Figs. 2B, 2D). CD11b- or F480-positive cell frequency among CD45^+^ cells did not differ in tumors from 4M and 12M mice (extended data Fig. 2I). Overall, these data define molecular changes occurring in macrophages of aged mice and highlight changes in select macrophage subsets that are either enriched or lost in the aged population.

#### Tumors in aging mice show an increase in the subpopulation of exhausted Cd8^+^ T cells expressing Gzmk

To determine whether increased numbers of immunosuppressor macrophages seen in tumors from older mice are linked with decreased cytotoxic T cell activity, we subjected the CD8^+^ T cell population in those samples to re-clustering, resulting in 12 distinct CD8^+^ T cell clusters (Fig. 3A; extended data Table 12). Among them, effector T cells (expressing high levels of Cd44, Cd28, Gzma, Gzmb, and Ccl5) were enriched in cluster T6 and effector CD8^+^ T cells expressing high levels of Ifng and Tnf were represented in cluster T7 (Fig. 3B). Cluster T9 represented exhausted T cells and showed the highest expression of Tox, Pdcd1, Gzmk, and Lag3, and cluster T8 was characterized by proliferating Cd8^+^ T cells expressing markers of exhaustion (Fig. 3B). The most robust age-related changes in Cd8^+^ T cell clusters seen in tumors from 4M, 12M, and 20M mice were identified as decreased effector Cd8^+^ cells (cluster T6) and increased proliferating and exhausted Cd8^+^ T cells (cluster T8) seen in 12M and 20M mice relative to 4M mice (Fig. 3C).

**Figure 3.**
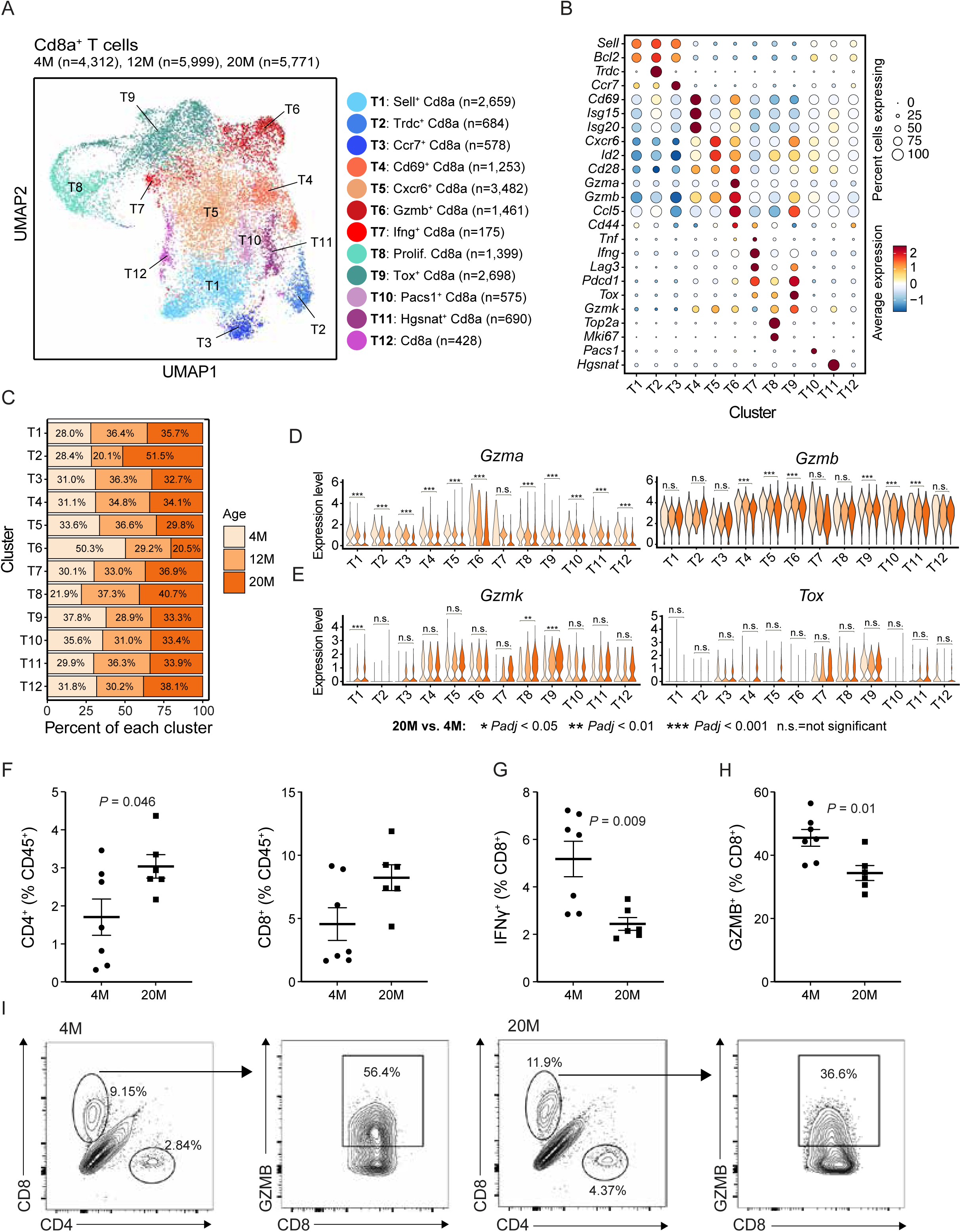
Tumors in aging mice show increases in a subpopulation of exhausted Gzmk-positive Cd8^+^ T cells. CD8^+^ cells (clusters C1-C3 from global UMAP in Figure 1D) from scRNA-seq analysis of CD45^+^ cells were re-clustered to obtain higher resolution clustering of CD8^+^ cells. (**A**) Integrated UMAP plot of CD8^+^ cells from 4M (n= 4,312 cells), 12M (n= 5,999 cells) and 20M (n= 5,771 cells) tumors showing 12 distinct clusters. (**B**) Dot plots showing scaled expression of marker genes used to identify CD8^+^ cell sub-clusters. (**C**) Bar graph showing the percentages of various sub-clusters of CD8^+^ cells for each cluster from 4M, 12M and 20M old mice. (**D-E**) Violin plots comparing expression of specific genes in CD8^+^ cell sub-clusters analyzed from 4M, 12M, and 20M mice. (**F**) Frequencies of CD4^+^ or CD8^+^ among CD45^+^ cells of tumors from 4M and 20M old mice. (**G-H**) Frequencies of IFNγ^+^ (G) and GZMB^+^ (H) among CD8^+^ cells in tumors from 4M and 20M old mice. (**I**) Representative flow cytometry plots showing GZMB^+^ expression by a gated subpopulation (CD8^+^ cells) of CD45.2^+^ cells from 4M and 20M old mice. n = 7 for 4M and n = 6 for 20M for panels F-H. Data in panel F-H were analyzed by unpaired *t*-test. Statistical significance is reported for 20M relative to 4M, computed using *FindMarkers* function in Seurat.

Assessment of genes implicated in cytolytic function across the 3 age groups revealed a notable decrease in Gzma and Gzmb expression in 20M compared to 4M samples of several T cell clusters (Fig. 3D). Also, relative to 4M mice, we observed a decrease in Ifng expression, albeit not significant in most 20M mice clusters (extended data Fig. 3A), in aged mice. We also identified a significant increase in the expression of Gzmk (an age-associated gene^26^) in clusters T8 and T9 of aged relative to young mice (Fig. 3E). Notably, age-associated increases in Gzmk expression were limited to clusters of exhausted, Tox-positive T cells (Fig. 3E), suggesting that Gzmk plays an age-dependent role in these phenotypes.

FACS analyses confirmed significant downregulation of IFNγ and Gzmb expression in CD8^+^ cells from tumors of 20M relative to 4M mice (Figs. 3F-3I). While the frequency of CD8^+^ among CD45^+^ cells was comparable in tumors from young and aged mice, we observed a significant increase in the frequency of CD4^+^ cells relative to CD45^+^ cells from tumors of 20M relative to 4M mice (Fig. 3F). When we analyzed tumors in 4M and 12M mice collected 12 days after YUMM1.7 cell injection we observed a decreased frequency of CD69^+^ effector cells among CD8^+^ and CD4^+^ cells in tumors from 12M relative to 4M mice (extended data Fig. 3B). Overall, these data show that CD8^+^ T cells seen in aging mice show decreased cytotoxic function, and exhausted T cells seen in aging mice are characterized by increased Gzmk expression.

#### Exhausted CD8^+^ T cells seen in melanoma tumors from patients show increased GZMK expression

To confirm these observations in human clinical specimens, we profiled gene expression in immune cells infiltrating tumors from two groups of melanoma patients representing younger and older populations, namely, ≤60 and ≥70 years old, respectively, using a previously published scRNA-seq dataset^27^. Immune cell clustering identified 12 clusters (Fig. 4A), each exhibiting a distinct gene expression pattern (Figs. 4B and 4C; extended data Table 13), including GNLY^+^ NK cells (cluster H7), CD3D^+^ T cells (clusters H1-H6), CSFR1^+^ myeloid cells (clusters H10-H11), FLT3^+^ DCs (cluster H12), and CD19^+^ B cells (clusters H8-H9). T-cell populations were distributed into cytotoxic CD8^+^ T cells expressing high levels of IFNG (cluster H2), exhausted CD8^+^ T cells expressing high levels of TOX (cluster H1), activated CD8^+^ T cells expressing CD44 (Cluster H3), TRDC^+^ T cells (cluster H4), TCF7^+^ T cells (cluster H5) and FOXP3^+^ T cells (cluster H6). Assessment of two patients age groups (≤ 60 and ≥ 70 years old), revealed that aging markedly decreased the frequency of T cells cluster H2, representing cytotoxic T cells expressing IFNG (Fig. 4D). Of note, we observed the most significant age-associated increase in GZMK expression in the exhausted sub-cluster of CD8^+^ T cells (Cluster H1), from pre-therapy patients (Fig. 4E). These observations corroborate our findings in mouse studies and identify a new subset of GZMK-positive exhausted T cells enriched in tumors from aging melanoma patients.

**Figure 4.**
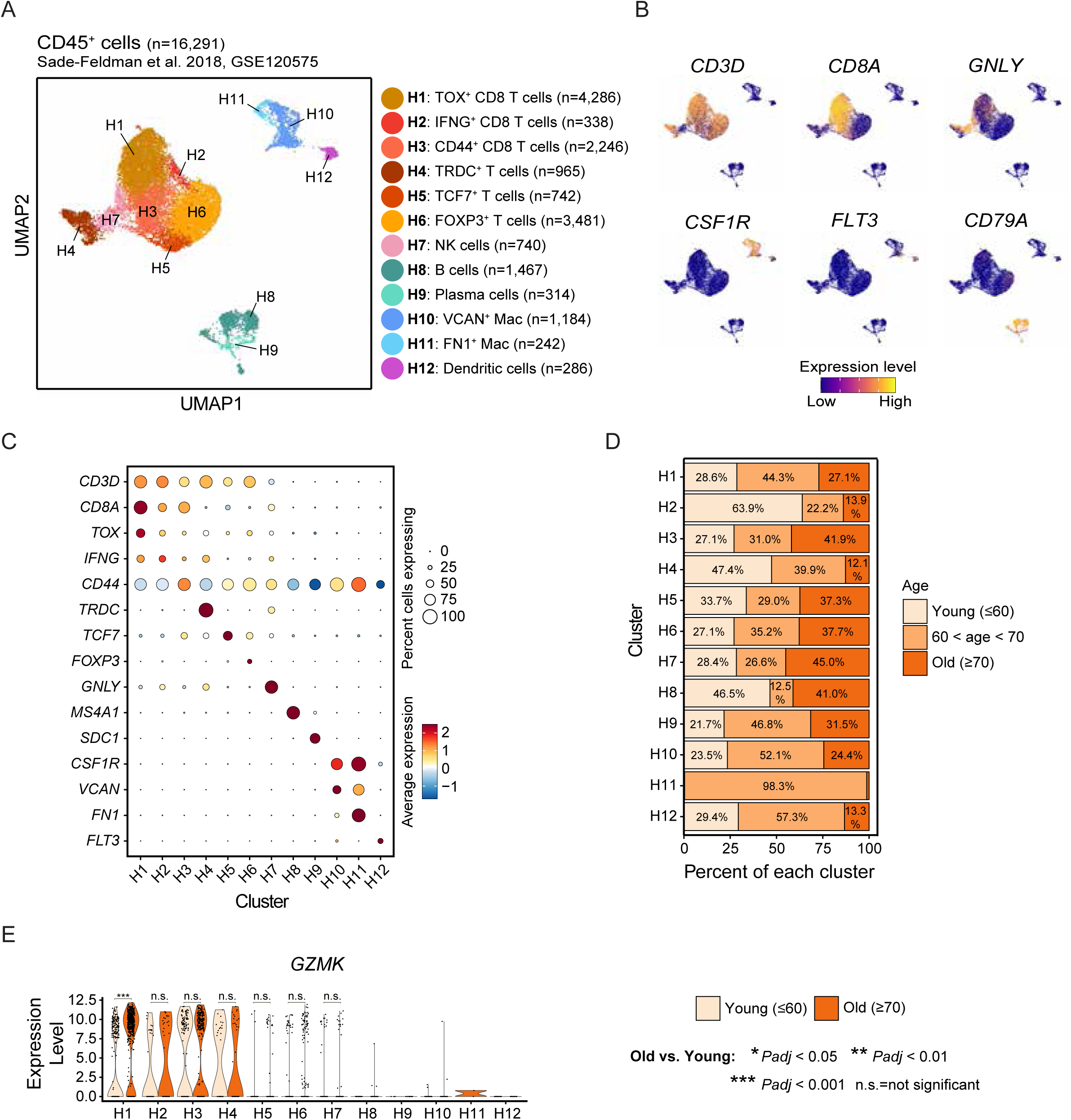
Exhausted CD8^+^ T cells from aged melanoma patients show increased GZMK expression. Immune cell clusters from scRNA-seq analysis of samples from melanoma patients (GSE120575) were stratified by age (≤ 60 compared to ≥ 70). (**A**) Integrated UMAP plot of CD45^+^ cells from melanoma patients showing 12 distinct clusters. Immune cell types identified using expression of specific cell markers are labeled. (**B**) UMAP plots showing expression of specific genes used to identify immune cell populations. (**C**) Dot plots showing scaled expression of marker genes used to identify immune cells populations. (**D**) Bar graph showing the percentage of various immune cells for each cluster based on analysis of melanoma patients in indicated age groups. (**E**) Violin plots comparing pre-therapy expression of GZMK in each immune cells cluster based on analysis of melanoma patients in indicated age groups. Statistical significance is reported for old (≥ 70) relative to young (≤ 60), computed using *FindMarkers* function in Seurat.

#### Immunogenic profiles exhibited by macrophages from aging mice are conserved in aging melanoma patients

To further assess the clinical relevance of our findings in mice, we performed sub-clustering of CSFR1-positive macrophages derived from tumors of melanoma patients stratified by age (as described above). Sub-clustering analysis (Figs. 5A, 5B; extended data Table 14) identified populations of macrophages similar to those identified in our mouse model, including TREM2- and APOE-expressing macrophages (cluster S4) that were represented in clusters M8 and M9 of mice-derived aged associated macrophages (Figs. 2C, 2D). Likewise, macrophages highly expressing TREM1 and FN1 (cluster S2) and interferon response genes (cluster S3) resembled mouse macrophage cluster M2, enriched in tumors from 4M mice (Figs. 2C, 2D). Furthermore, we observed macrophages exhibiting high expression of S100A8 and S100A9 (cluster S1) and a cluster expressing genes involved in lipid metabolism (such as CD36; cluster S5). Overall, our classification of macrophage clusters from melanoma patients (Figs. 5A, 5B) resembled that seen in melanoma-bearing mice (Figs. 2A, 2B). Of note, we observed enrichment of the population of APOE/TREM2-positive macrophages (cluster S4) in tumors from aging melanoma patients (≥ 70 years old), while macrophages exhibiting the interferon response gene signature (cluster S3) were enriched in tumors from younger patients (≤ 60 years old; Fig. 5C), which were also patterns seen in our mouse melanoma model.

**Figure 5.**
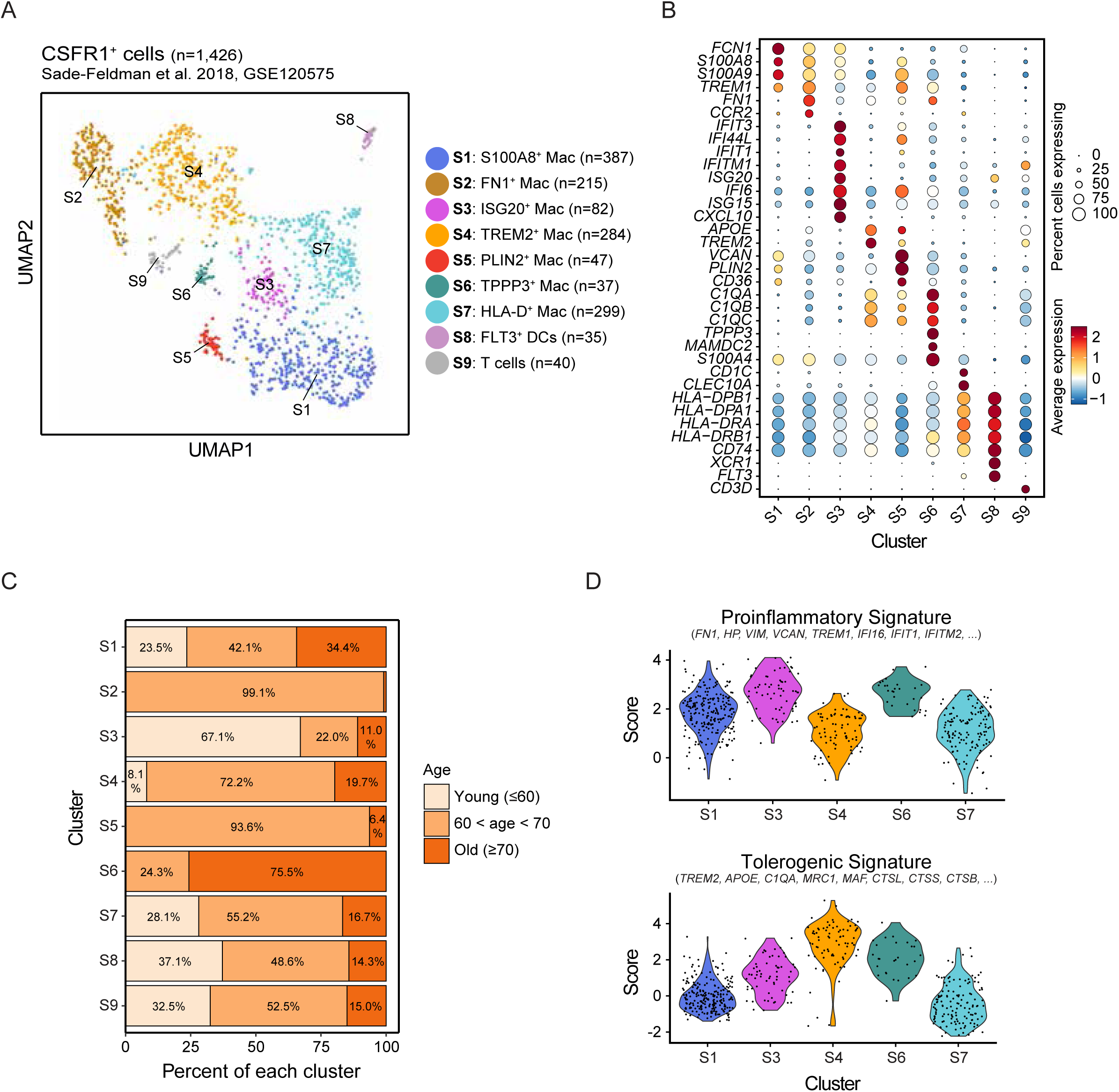
Immunogenic profiles exhibited by macrophages from aging mice are conserved in aging melanoma patients. Myeloid cell clusters (H10 and H11 from global UMAP in Figure 4A) from human patient scRNA-seq analysis of CD45^+^ cells were re-clustered to obtain higher resolution clustering of myeloid cells. (**A**) Integrated UMAP plot of myeloid cells from < 60 (n=279 cells), >60 and <70 (n=855 cells) and >70 (n=292 cells) years old melanoma patients showing 9 distinct clusters. **(B**) Dot plots showing scaled expression of marker genes used to identify specific myeloid cell sub-clusters. (**C**) Bar graph showing the percentage of various subclusters of macrophages from melanoma patients ≤ 60 and ≥ 70 years old. (**D-E**) Violin plots comparing tolerogenic (D) and inflammatory (E) signature score in each macrophages’ cluster (only clusters represented in ≤ 60 and ≥ 70 years old melanoma patients are shown).

Analysis of markers expressed explicitly in mouse cluster M2, which is predominant in 4M mice (extended data Tables 2, 15), revealed an inflammatory gene signature. Likewise, analysis of markers specific to mouse cluster M8, which is more prominent in 20M mice, indicated a tolerogenic gene signature (extended data Tables 2, 15). When we evaluated both signatures in macrophage clusters from melanoma patients stratified by age (≤ 60, and ≥ 70), we observed an increase in the tolerogenic signature in cluster S4, which is more represented in cells from tumors from aged melanoma patients (≥ 70 years old) that also express TREM2 and APOE (Fig. 5D). The inflammatory signature was enriched in cluster S3, which is more represented in the younger melanoma patient population (≤ 60 years old) and expresses interferon response genes (Fig. 5E). The relatively small number of cells expressing TREM1 and FN1 (cluster S2) in patients ≤ 60 and ≥ 70 years old, which was enriched in 4M mouse macrophages (cluster M2), prevented us from comparing human and mouse data (Figs. 2A-D). Nonetheless, using the TCGA dataset as a resource, we analyzed TREM1 expression in skin cutaneous melanoma (SKCM) samples stratified by age and identified a significant increase in TREM1 expression in the younger melanoma patient group (extended data Fig. 4A). Analysis of an independent data set^28^ confirmed age-associated TREM1 expression (extended data Fig. 4B). Although analysis of TREM2 expression using the TCGA dataset of SKCM samples did not reveal a significant difference between the two age groups (likely due to a different distribution of macrophage subpopulations in metastatic vs. primary tumors), previous published data of lung adenocarcinoma (LUAD), breast cancer (BRCA), prostate adenocarcinoma (PRAD), sarcoma (SARC) and thymoma (THYM) reported a significantly higher expression of TREM2 in tumor specimens from the aged specimens, when compared to those obtained from the younger patients^29^. Thus, age-associated increases in the number of TREM2-expressing macrophages were observed in melanoma tumors from aged mice and in different tumor types found in older patients.

#### TREM1 and TREM2 determine age-associated melanoma progression and metastasis

To assess how Trem1- and Trem2-expressing macrophages impact melanoma growth and its metastasis to lung, we inhibited TREM1 or TREM2 activity in 4M and 20M melanoma-bearing mice. To block TREM1 activity, we treated mice with the TREM1 pharmacological inhibitor VJDT 4 days after YUMM1.7 melanoma cell inoculation and repeated treatment every other day until tumors were collected. Of note, the YUMM1.7 melanoma cell line harbors a low number of somatic genetic changes and is minimally immunogenic, making it largely resistant to PD1 therapy^30^. Pharmacological TREM1 inhibition delayed the growth of anti-PD1-resistant melanoma cells in 4M but not 20M mice (Fig. 6A), consistent with our observation that younger mice have a higher representation of Trem1-expressing macrophages.

**Figure 6.**
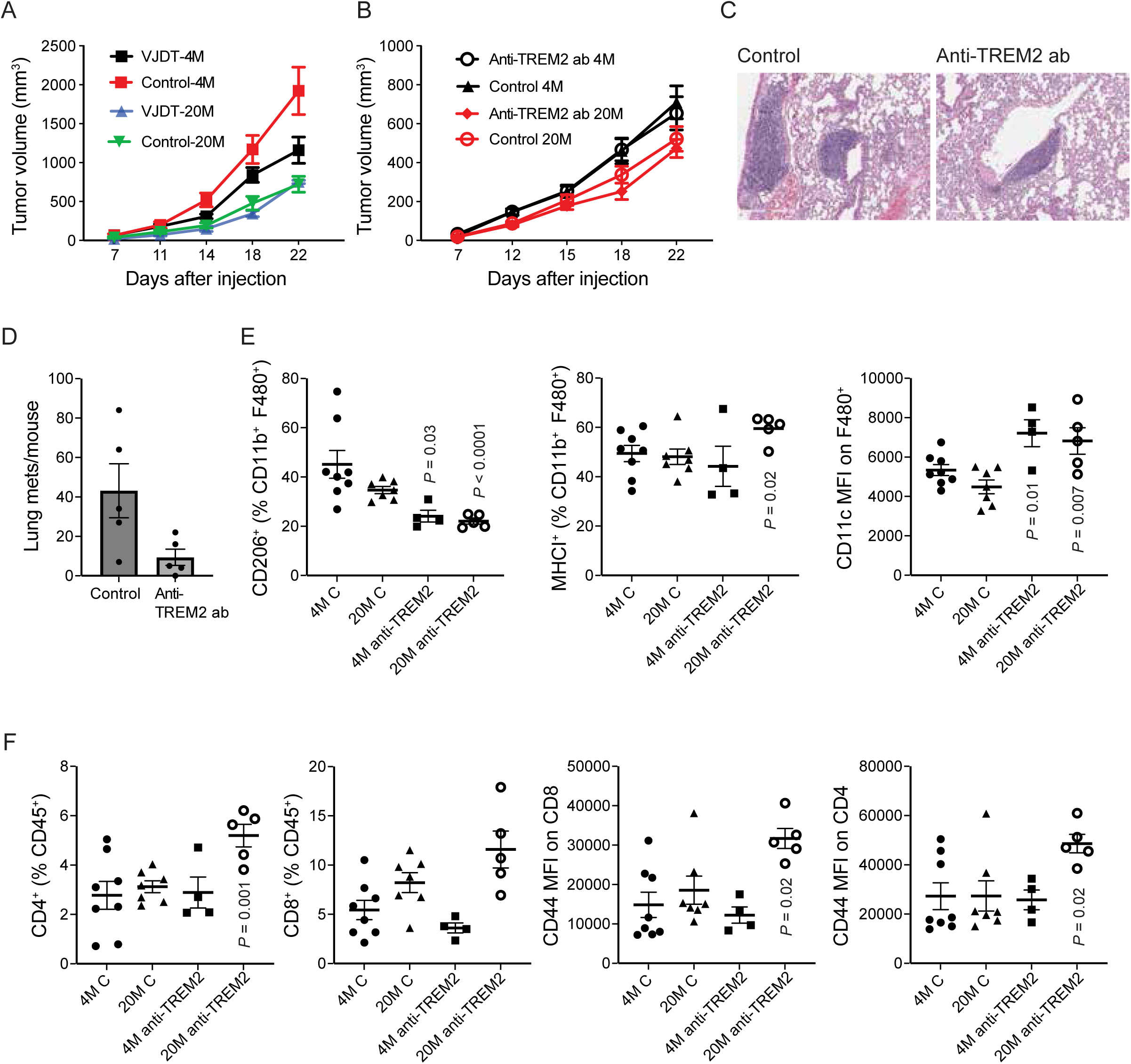
Function of TREM1 and TREM2 in age-associated melanoma progression and metastasis. (**A-B**) YUMM1.7 melanoma cells (500,000) were injected s.c. into the flanks of 4M (young), and 20 months (old). Four days later mice were injected i.p. with either TREM1 inhibitor VJDT or vehicle (control) (A) and injections were repeated every other day until day 20. A separate group of melanoma-bearing mice were treated with anti-TREM2 antibody (B) starting at day 5 after melanoma cell injection and repeated every three days for a total of 4 treatments. Isotype control served as control (B). n = 5 for 4M control and TREM1 treated, n = 4 for 20M in panel A; n = 8 for 4M control, n = 5 for 4M TREM2 treated, n = 7 for 20M control, n = 5 for 20M TREM2 treated in panel B. (**C**) Representative H&E staining of lung tissues from 20M melanoma tumor-bearing mice treated either with isotype control (left) or anti-TREM2 antibody (right). Lung tissues were analyzed 22 days after melanoma cell injection. (**D**) Quantification of lung metastases following treatment of mice with control or anti-TREM2 antibody as described in (B). Tissues were subjected to IHC staining for S100 to detect metastatic melanoma cells (more than 10 S100+ cells per lesion). n = 5 for each group. Scale bar, 300 μM. (**E**) Frequencies of CD206^+^ and MHCI^+^ among CD11b^+^ and F480^+^ cells, and expression of CD11c^+^ cells on F480^+^ cells of tumors from 4M and 20M old mice treated with isotype control or anti-TREM2 antibodies. (**F**) Frequencies of CD4^+^ and CD8^+^ among CD45.2^+^ cells, and expression of CD44^+^ cells on CD4^+^ or CD8^+^ cells of tumors from 4M and 20M old mice treated with isotype control or anti-TREM2 antibodies. n = 8 for 4M control, n = 4 for 4M TREM2 treated, n = 7 for 20M control, n = 5 for 20M TREM2 treated in panels E and F. Data in panels A, B, D, E and F are presented as means ± SEM. Data in panel A were analyzed by two-way ANOVA. Data in panel D, E and F were analyzed by unpaired *t*-test and compared to controls of the same age group.

To inhibit TREM2, we injected 4M and 20M melanoma-bearing mice with anti-TREM2-neutralizing antibodies^22,31^. Anti-TREM2 treatment did not alter tumor growth over time in either age group (Fig. 6B); however, anti-TREM2 antibody injection significantly inhibited lung metastasis in 20M mice (Figs. 6C, 6D, extended data Figs. 5A, 5B), suggesting that TREM2 blockade is effective in inhibiting lung metastasis of poorly immunogenic melanomas.

We next asked if TREM2 inhibition altered phenotypes of immune cells infiltrating tumors in 4M and 20M mice. Based on FACS analysis, we observed that in the presence of TREM2 inhibition, there was a significant decrease in the M2 marker CD206 among CD11b^+^ and F480^+^ cells in mice treated with anti-TREM2 antibody and independent of age (Fig. 6E), while the frequency of CD11b- and F480-expressing cells among CD45^+^ cells was comparable independent of age and treatment (extended data Fig. 5C), suggesting that macrophages become polarized to an M1-like phenotype. Relative to untreated controls, we also observed upregulation of CD11c (Fig. 6C), a pro-phagocytic integrin induced in inflammatory macrophages and implicated in anti-tumor immunity, in macrophages subjected to TREM2 inhibition in 4M and 20M mice. Of note, anti-TREM2 neutralizing antibody was more efficient in inducing MHCI expression in macrophages from tumors of 20M relative to 4M mice. Notably, TREM2 inhibition increased infiltration and activity of T cells in 20M relative to 4M mice. Anti-TREM2 antibodies significantly increased the number of CD4^+^ cells in 20M mice, and increased, albeit not significantly, the number of CD8^+^ cells (Fig. 6F). Of interest, T cells were significantly activated in 20M mice, based on increased frequency of CD44^+^ cells among CD4^+^ and CD8^+^ cells, after anti-TREM2 treatment (Fig. 6F). These data strongly suggest that TREM1 functions in melanoma progression in younger mice and that TREM2 promotes metastatic activity of melanoma in aged mice.

## Discussion

Immune cell function in melanoma progression is well studied, but how aging impacts tumor infiltration by immune cells remains poorly understood. Here, we show that aging enhances the development of an immunosuppressive tumor microenvironment likely by decreasing cytotoxic T cell function and polarizing macrophages to tolerogenic phenotypes both in melanoma-bearing mice and in patients with melanoma. These studies highlight selective and age-associated enrichment of TREM1- and TREM2-expressing macrophages during melanoma progression. Therapeutically, we show that TREM2 inhibition prevents lung metastasis in aged mice, while TREM1 inhibition hampers tumor growth in younger mice. TREMs, or Triggering Receptors Expressed on Myeloid cells, are a family of cell surface receptors that modulate the threshold and duration of myeloid cell responses^32,33^. TREM1 and TREM2 are best characterized and reportedly function in different diseases, including cancers^33^: TREM1 is well known for pro-inflammatory function^34,35^, while TREM2 plays an immunosuppressive role. Despite these opposing functions, both receptors promote tumor growth. TREM1 activity reportedly sustains inflammation and contributes to cancer progression^36^. High TREM1 levels are associated with poorer survival in non-small cell lung cancers^37^ and hepatocellular carcinoma, while TREM1 function in age-associated melanoma progression has not been reported.

Here we show enrichment of TAMs expressing TREM1 preferentially in younger relative to older populations. Using young and aged mice injected with a melanoma cell line (YUMM1.7), which is known to be resistant to immunotherapy^30^, we show that TREM1 blockade inhibits tumor growth in younger but not older mice. Moreover, using two independent datasets (TCGA and^28^), we confirmed increased TREM1 expression in tumor samples from younger versus older melanoma patients, corroborating findings in our mouse model. These observations suggest that aging switches melanoma-associated macrophages from an inflammatory to an immunosuppressor phenotype, and these outcomes could be exploited to improve immunotherapy efficacy based on age.

The TREM2 receptor can bind lipids, and its expression on macrophages is linked to the progression of Alzheimer’s disease, atherosclerosis, and metabolic disorders^32,38,39^. TREM2 is expressed on TAMs infiltrating tumors in both human cancers and mouse models, including sarcoma, colorectal, and breast cancers^40,41,31,42^. TREM2 genetic or pharmacological inactivation antagonizes the activity of immunosuppressive macrophages and increases CD8 T cell effector function, decreasing tumor growth in several cancers^43,40,44^. Here for the first time, we show age-associated enrichment of TREM2/APOE-expressing macrophages in a melanoma mouse model and establish a novel role for these cells in promoting lung metastasis. Indeed, while others have reported a role for TREM2 in solid tumors^45,41,31,46^, this study highlights the function of TREM2-expressing macrophages in metastasis. In melanoma patients, high Trem2 expression was previously associated with resistance to immunotherapy^47^, and a recent study has shown age-dependent regulation of TREM2 expression in several cancers^29^, corroborating our findings in melanoma infiltrated macrophages.

Other recent studies have described the effects of aging on tumor infiltration by T cells^48,49,50^. Here we confirm age-associated decreases in cytotoxic T cells in our mouse model and in melanoma patients and identify a subset of dysfunctional cytotoxic T cells expressing exhaustion markers enriched in melanoma tumors from aged mice. This T cell subset expresses high levels of Granzyme K, an observation made in mice and melanoma patients. Age-related increases in granzyme K expression in CD8^+^ cells were recently described in multiple organs in both mice and humans, and secreted Granzyme K is thought to promote inflammation^26^. Granzymes K, A, and B constitute a family of serine proteases mainly expressed by CD8^+^ T and NK cells that function primarily to induce cell death in virally infected cells and tumor cells. Granzyme K is structurally similar to Granzyme A, but each has its own function and substrates^51^. Of note, increased Granzyme K expression in CD8 cells was previously associated with poor prognosis in colorectal cancer patients^52^. It is currently not known whether secreted Granzyme K promotes an immunosuppressive function in T cells, and further studies are needed to determine if Granzyme K could be targeted to increase T cell toxicity against melanoma in the context of aging.

How aging enriches specific subsets of melanoma-associated macrophages remains unknown. Aging leads to declines in immune cell function and epigenetic regulation plays an important role in immune-senescence and age-associated inflammation^53,54,55^. Aging can promote the accumulation of cytokines, hormones, free fatty acid, and low-density lipoproteins, activating macrophages^56,57^. TREM2 binds to phospholipids and sulfatides ^58^, including APOE ^59,60^ and age-associated lipids in the tumor microenvironment may function in the enrichment of TREM2-expressing macrophages or enhance their function.

A limitation of this study is that in our melanoma mouse model, young mice do not exhibit lung metastasis. Therefore, we could not assess the potential impact of TREM2 inhibition on lung metastasis in young mice. While we identify a novel TREM2 function in promoting lung metastasis, our study does not address the contribution of TREM2-expressing macrophages from a primary tumor versus a potential metastatic function of TREM2-expressing macrophages found in the lung.

Overall, these studies show that aging is a factor in establishing an immunosuppressive tumor microenvironment characterized by subsets of macrophages with a tolerogenic phenotype and by dysfunctional CD8^+^ cells with an exhausted phenotype. We show that macrophages expressing TREM1 and TREM2 are differentially enriched in an age-dependent fashion in both a mouse model and melanoma patients. Finally, we demonstrate that TREM1 inhibition decreases melanoma growth in young but not old mice and that TREM2 inhibition antagonizes lung metastasis. These data could be relevant in identifying novel targets to improve immunotherapy based on the age of melanoma patients.

## Online Methods

### Mouse tumor models and ethics

All experimental animal procedures were approved by the Institutional Animal Care and Use Committee (IACUC) of Sanford Burnham Prebys Medical Discovery Institute. The mice were housed under a 12-h light–dark cycle at controlled ambient temperature (22 ± 1 °C) and relative humidity (40–70%). Adequate standard chow and water were available ad libitum via cage lids. All mice were on a C57BL/6 genetic background. Aged mice were obtained from the NIA repository. Male mice were used to obtain animal data in this study and are representative of at least three independent cohorts. *Braf^V^*^600^*^E/+^; Pten^−/−^; Cdkn2a^−/−^* mouse melanoma cells (YUMM1.7) were kindly provided by Dr. Marcus Bosenberg. For tumor growth experiments, mice were injected subcutaneously (s.c.) with 500,000 YUMM1.7 cells. Tumor volumes were measured 3 times a week. Tumors were collected 22 days after inoculation, unless otherwise noted.

### In vivo treatments

Anti-TREM2 neutralizing antibody was purchased from Leinco Technologies. Mice were injected intraperitoneally (i.p.) with anti-TREM2 antibody (Leinco Technologies, clone 178; 10 mg/kg mouse) starting on day 5 after melanoma injection and every three days for a total of 4 treatments as previously reported^22,31^. Anti-hILT1 (Leinco Technologies, clone 135.5) was used as control isotype^22,31^. For pharmacological inhibition of TREM1, 20mg/kg of VJDT (MedChemexpress) or vehicle (DMSO), were administered (i.p.) in mice at day 4 after melanoma cells injection and continued alternate days util day 20 as previously reported^36^.

### Tissue digestion

Tissues were excised, minced, and digested with 1 mg/ml collagenase D (Roche) and 100 µg/ml DNase I (Sigma) at 37°C for 45 minutes. Digests were then passed through a 70-μm cell strainer to generate a single-cell suspension. Cells were washed twice with PBS containing 2 mM EDTA and stained for flow cytometry.

### Flow cytometry

Tissue-derived single-cell suspensions were washed twice with FACS staining buffer, fixed 15 min with 1% formaldehyde in PBS, washed twice, and resuspended in FACS staining buffer for staining with specific antibodies, followed by fixation with 1% formaldehyde and then flow cytometry analysis. The following antibodies were purchased from Biolegend: CD45.2 (104), CD8a (53-6.7), CD4 (GK1.5), CD11c (N418), CD11b (M1/70), MHC class I (AF6-88.5), CD206 (C068C2), F4/80 (BM8), CD44 (IM7),CD69 (QA17A41), IFNg (XMG1.2), GRANZYME B (GB11) was purchased from BD Biosciences, TREM2 (ab313953) was purchased from Abcam and TREM1 (747899) was purchased from R&D. All data were collected on an LSRFortessa cell analyzer (BD Biosciences) and analyzed using FlowJo Software (Tree Star).

### Histology and immunofluorescence on tissue

Lungs were fixed in 4% formalin overnight at 4°C, washed with PBS, paraffin-embedded, cut into 5 μm-thick sections, and stained with H&E. For immunofluorescence, sections were deparaffinized, rehydrated and washed in PBS. Antigen retrieval was performed in a pressure cooker (Decloaking chamber, Biocare Medical) in citrate buffer (pH 6.0). Ki67 (AbCam Ab15580) or S100 (Thermo-Fisher 500-2144) immunostaining was performed by incubating sections overnight at 4°C with antibodies in Dako antibody diluent. Alexa Fluor 594-conjugated secondary antibodies was added for 1h at room temperature (Molecular Probes), and nuclei were counterstained using SlowFade Gold Antifade reagent (Vector) with 4′,6-diamidino-2-phenylindole (DAPI, Vector). Image data were obtained using Olympus TH4–100 microscope and using Slidebook 4.1 digital microscopy.

### Quantification of lung metastasis

Fixed and embedded lung tissues were sliced to obtain five serial sections per lung. H&E and S100 antibody (used as a marker for melanoma cells^61^ staining were used to quantify number of nodules S100 positive (10 or more cells) using Olympus TH4–100 microscope.

### Single-cell library preparation and sequencing

4-, 12-, and 20M old mice (7 mice for each age) were sacrificed 18 days after tumor cell inoculation. Tumors were digested as described above. Single cell suspensions were washed with 1X PBS, 4% FBS before incubation for 20 min on ice at 5×10^7^ cells/ml with 500 ng/ml Fc block (2.4G2, BD Pharmingen). Cells were then incubated for 1 hour on ice with AF700 conjugated CD45.2 monoclonal antibody (104, Biolegend). For scRNA-seq libraries, DAPI negative (live) CD45^+^ cells were sorted using a flow cytometer, and sorted cells were resuspended in RPMI for counting. Libraries were prepared using the Fluent Biosciences PIP-seq v4.0 Plus T20 3’ Single Cell RNA kit according to the User Guide. Libraries were sequenced on an Element Biosciences AVITI sequencer with 60×100bp sequencing to a read depth of 20k – 27k reads per cell.

### PIP-seq data pre-processing and analysis

PIP-seq samples were processed using Pipseeker pipeline version 2.1.3, STAR aligner version 2.7.10a^62^, mouse genome version GRCm38, and Gencode annotations version M25^63^. Pipseeker pipeline outputs for 4M (sensitivity = 2), 12M (sensitivity = 3), and 20M (sensitivity = 4) were used for integration and analysis. Gene count matrices for 4, 12, 20M old mouse PIP-seq samples were processed using Seurat^64^ version 4.0.5 and R version 4.0.2. Count matrix for each sample was converted to Seurat object by retaining genes expressed in minimum of 5 cells and cells expressing a minimum of 200 genes. Cells expressing >10% mitochondrial genes (to remove dead/low quality cells) and top 2% genes expressed (to remove potential doublets/multiplets) were discarded. Seurat objects for 4M, 12M, and 24M mouse samples were merged and integration of samples was performed using *sctransform* normalization method (https://satijalab.org/seurat/articles/integration_introduction.html#performing-integration-on-datasets-normalized-with-sctransform-1) by regressing out “nCount_RNA” and percentage mitochondrial content. Top 3,000 variable genes were used in *SelectIntegrationFeatures* step. The three datasets were integrated using *FindIntegrationAnchors(normalization.method = "SCT")* and *IntegrateData(normalization.method = "SCT").* PCA components were computed using *RunPCA().* Cell Clusters were computed using *RunUMAP(dims=1:30)*, *FindNeighbors*, and *FindClusters (resolution=0.5)* resulting in 14 cell clusters. For visualization of gene expression and differential expression analysis, default assay was set to *“RNA”* and gene counts normalized using *NormalizaData*. Cluster markers were found using *FindAllMarkers.* Differential expression analysis comparisons were performed using *FindMarkers (test.use = “MAST”)*. Plots were prepared using Seurat and ggplot2^65^. Pathway analyses were performed in Ingenuity Pathway Analysis (Qiagen, Redwood City, USA). Main and supplemental plots were prepared using ggplot2^68^. Complex Heatmap, and Seurat Re-clustering of myeloid and T cell clusters were performed by selecting cells from these cell types and repeating the above-described method. For reclustering, 2000 highest variably expressed genes were selected for myeloid and T cell reclustering.

### Integrated analysis of scRNA-seq data from GSE120575

Normalized gene expression (TPM), cell annotation, and sample information were obtained from the GEO record (GSE120575)^27^. TPM data was converted to Seurat object using *CreateSeuratObject*. 2000 highest variably expressed genes were selected. Data was scaled using *ScaleData* and linear dimension reduction performed using *RunPCA*. Cell Clusters were computed using *RunUMAP (reduction=”harmony”, dims=1:15)*, *FindNeighbors (reduction=”harmony”, dims=1:15)*, and *FindClusters (resolution=0.5)*, resulting in 12 cell clusters. Cluster markers were identified using *FindAllMarkers.* Differential expression analysis comparisons were performed using *FindMarkers().* Average expression of proinflammatory and tolerogenic gene signatures^35^ (extended data Table 15) were computed using *AddModuleScore*.

### Analysis from TCGA and GEO data

For TCGA-SKCM data^66^, melanoma patient samples on cBioPortal^67^ were stratified according to age and patients ≤60 years old or > 70 years old were used for TREM1 expression comparison. P-value was computed using Student’s t-test.

Normalized gene expression (FPKM) for patients from published dataset^28^ was downloaded from GEO under accession GSE78220. Patients ≤ 60 years old were labelled as young while patients ≥70 years old were labelled as old. TREM1 expression was compared between young and old patient groups. P-value was computed using one-sided Student’s t-test.

### Statistical Analysis

GraphPad Prism version 10 was used for statistical analysis. Differences between two groups were assessed using two-tailed unpaired *t* test. Two-way ANOVA with Bonferroni’s multiple comparison test was used to evaluate experiments involving multiple groups. Differentially expressed genes in scRNA-seq clusters were identified using *FindMarkers()* and *MAST* test in Seurat. Survival analyses were computed using log-rank test.

## Data availability

All data needed to evaluate the conclusions in the paper are present in the paper and/or the Supplementary Materials. Additional information may be requested from the corresponding author. PIP-seq raw and processed data have been deposited in GEO (GSE280640).

## ACKNOWLEDGMENTS

We thank members of the Ronai lab for discussions. We thank Dr. Marcus Bosenberg (Yale University) for providing the YUMM1.7 melanoma cells. We also thank shared resources at the SBP Cancer Center supported by NCI grant P30CA030199. Support by NCI grant R35CA197465 (to ZR) and NIA support grants P01 AG073084 (PA), and R00AG068303 (XT) are gratefully acknowledged.

## AUTHOR CONTRIBUTIONS

MS and ZAR conceived the study; MS, PB, YF and RP performed experiments; MS, RM, KV, AT, XT, PA and ZAR analyzed data; MS, RM and ZAR wrote the manuscript.

## MATERIALS & CORRESPONDENCE

Correspondence and requests for materials should be addressed to Marzia Scortegagna, or to Ze’ev A. Ronai

## Extended data; Figures Legend

**Extended Figure 1.**
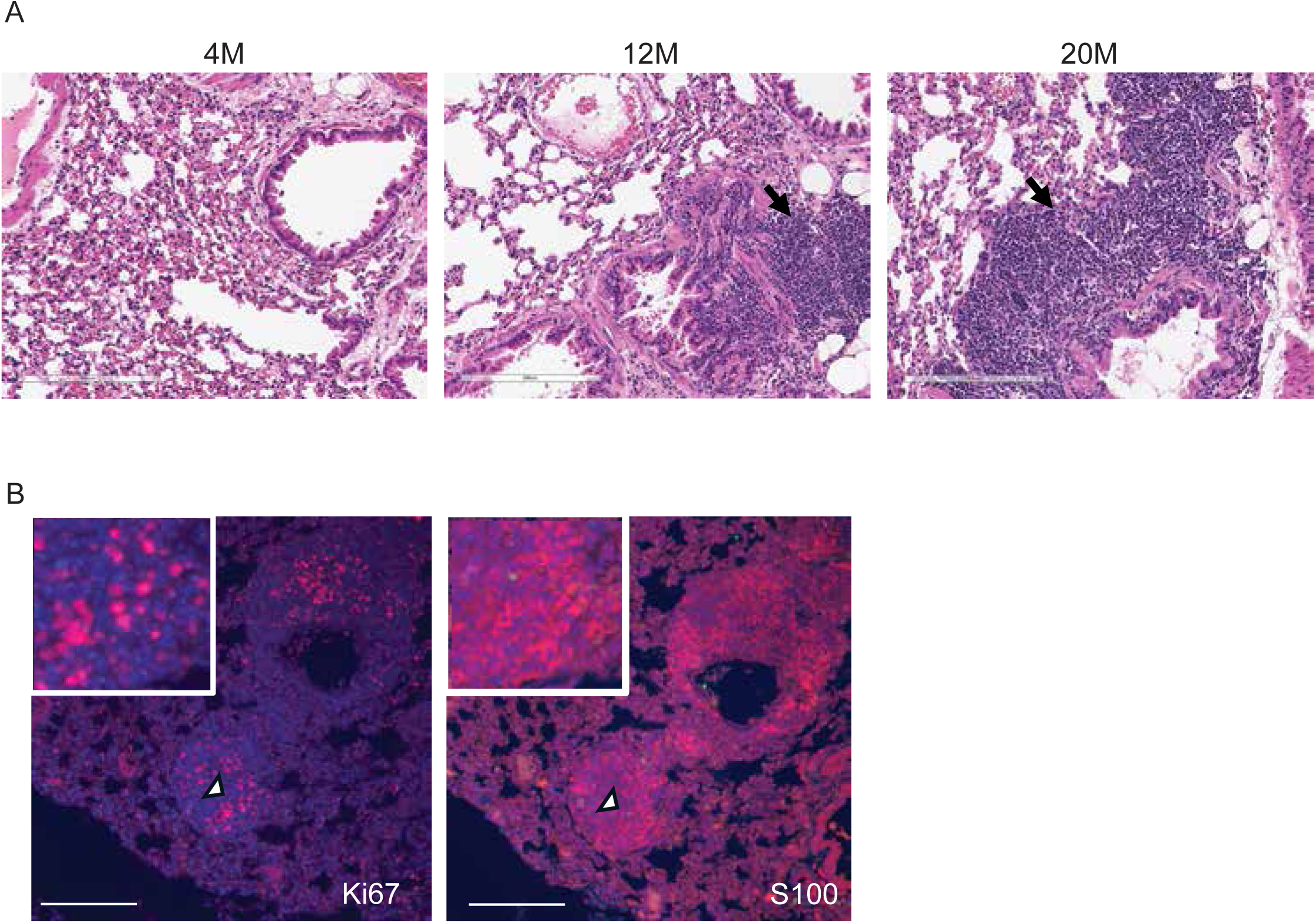
Lung metastasis increases with age in melanoma tumor-bearing mice. (**A, B**) YUMM1.7 melanoma cells (500,000) were injected s.c. into the flank of 4M, 12M and 20M old mice and 22 days later lungs were collected and processed to obtain 5 serial sections per lung. (A) H&E of lung tissues from indicated age groups. Scale bar, 200 μM. (B) IHC staining of either S100 or Ki67 to detect respective metastasis or cell proliferation. Scale bar, 100 μM.

**Extended Figure 2.**
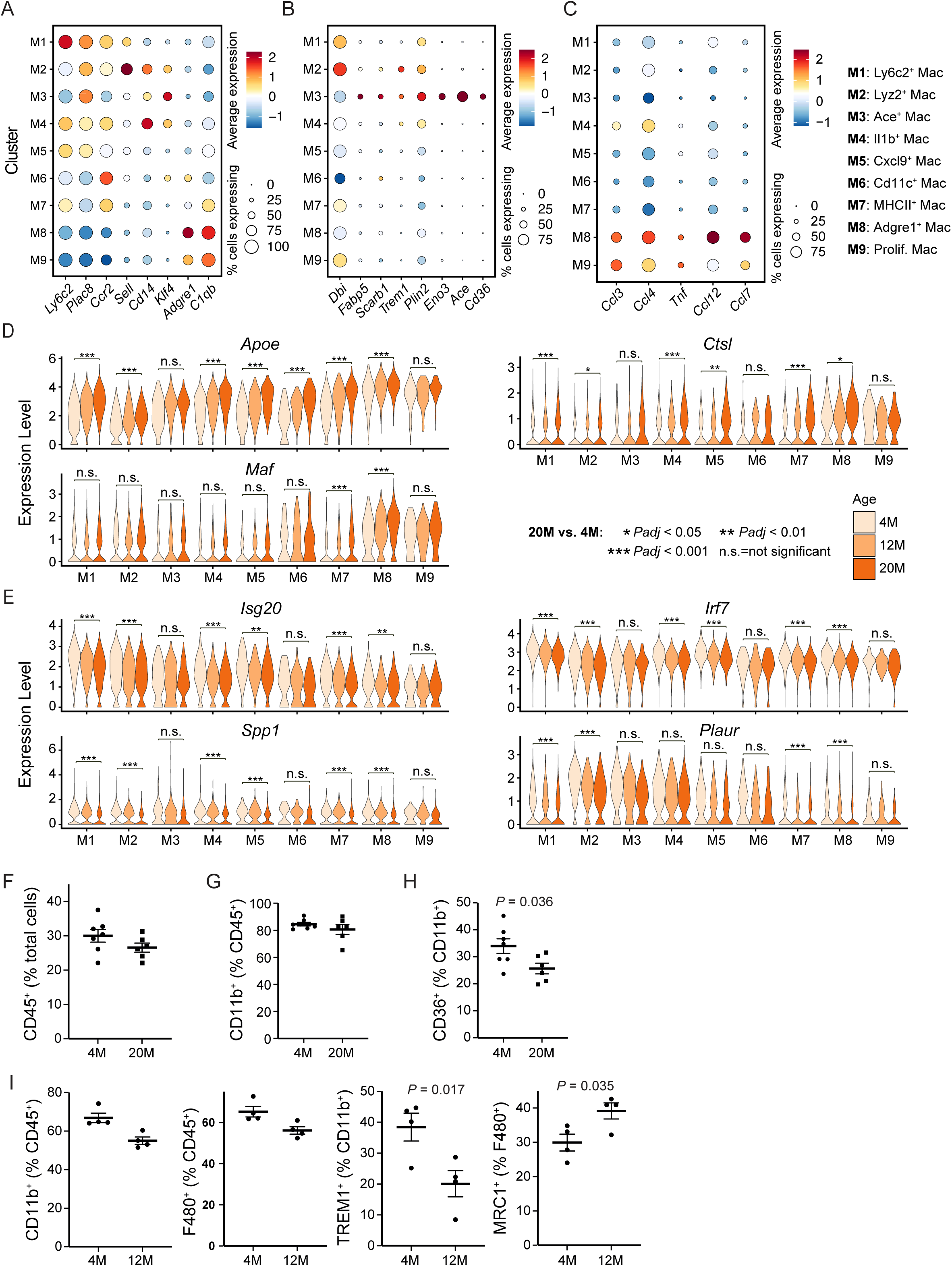
Aging alters macrophage gene expression patterns. Myeloid cell clusters (10 and 11 from global UMAP in Figure 4A) from scRNA-seq analysis of CD45^+^ cells were re-clustered to obtain higher resolution clustering of myeloid cells. (**A-C**) Dot plots showing scaled expression of marker genes used to identify monocyte markers (A), genes involved in lipid metabolism (B) and chemokines (C) in each sub-clusters of myeloid cells. (**D-E**) Violin plots comparing expression of specific genes in 4M, 12M, 20M old macrophages subclusters. Statistical significance is reported for 20M relative to 4M performed using *FindMarkers* in Seurat. (**F-H**) Frequencies of CD45.2^+^ among total cells (F), frequencies of CD11b^+^ among CD45.2^+^ cells (G) and frequencies of CD36^+^ among CD11b^+^ cells (H) in tumors from 4M and 20M old mice. (**I**) Frequencies of CD11b^+^ and F480^+^ among CD45.2^+^ cells, of TREM1^+^ among CD11b^+^ cells and of MRC1^+^ among F480^+^ cells in tumors from 4M and 12M old mice. n = 7 for 4M and n = 6 for 20M in panels F-H. n = 4 for 4M and 12M in panel I. Data in panels F-I were analyzed by unpaired *t*-test.

**Extended Figure 3.**
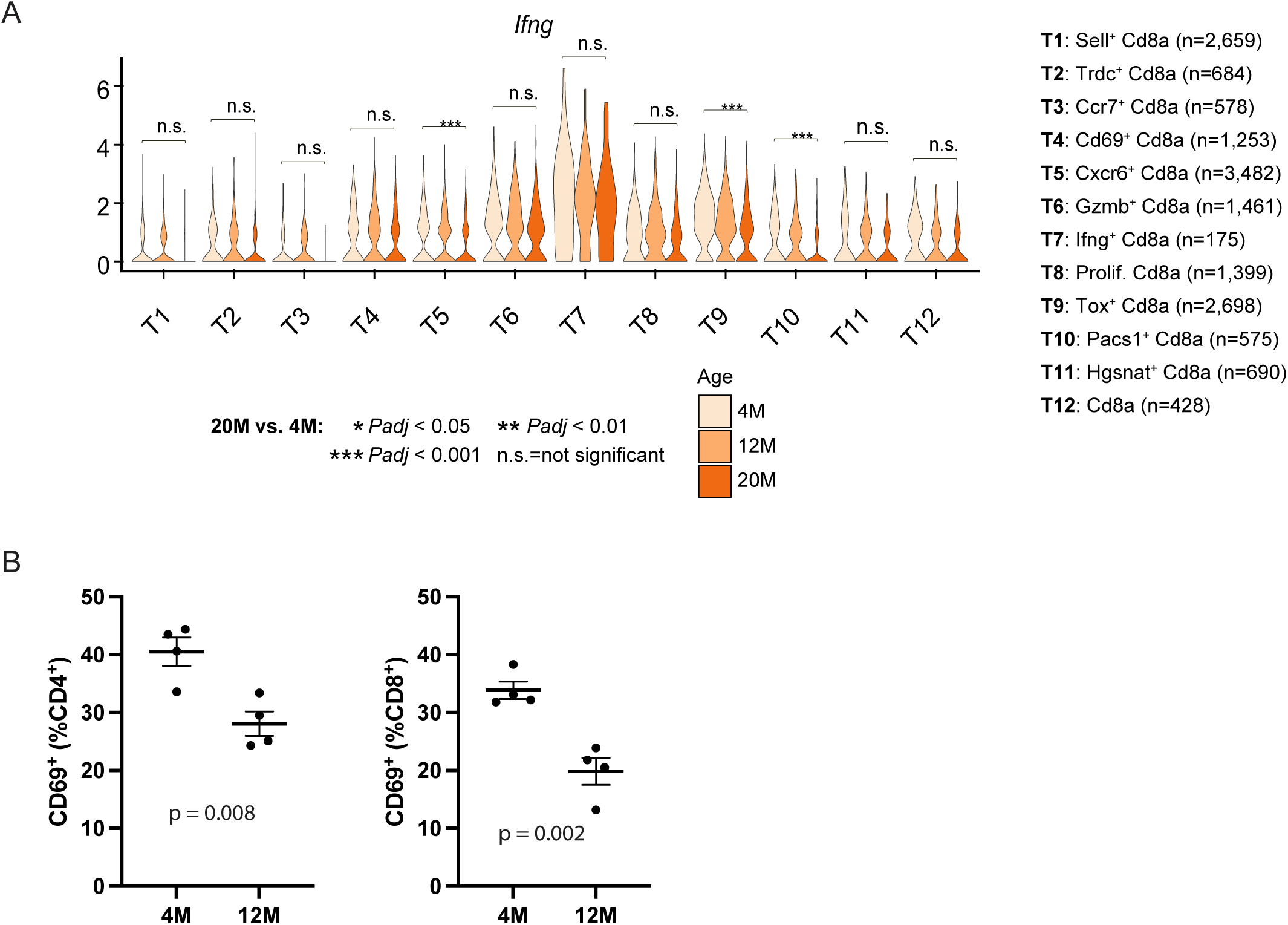
Aging alters gene expression in CD8^+^ T cells. (**A**) Violin plots comparing expression of specific genes in CD8^+^ cell subclusters from 4M, 12M, 20M mice. Statistical significance is reported for 20M relative to 4M performed using *FindMarkers* in Seurat. (**B**) Frequencies of CD69^+^ cells among CD4^+^ or CD8^+^ cells in tumors from 4M and 12M mice. N = 4 for 4M and 12M in panel B. Data in panel B were analyzed by unpaired *t*-test.

**Extended Figure 4.**
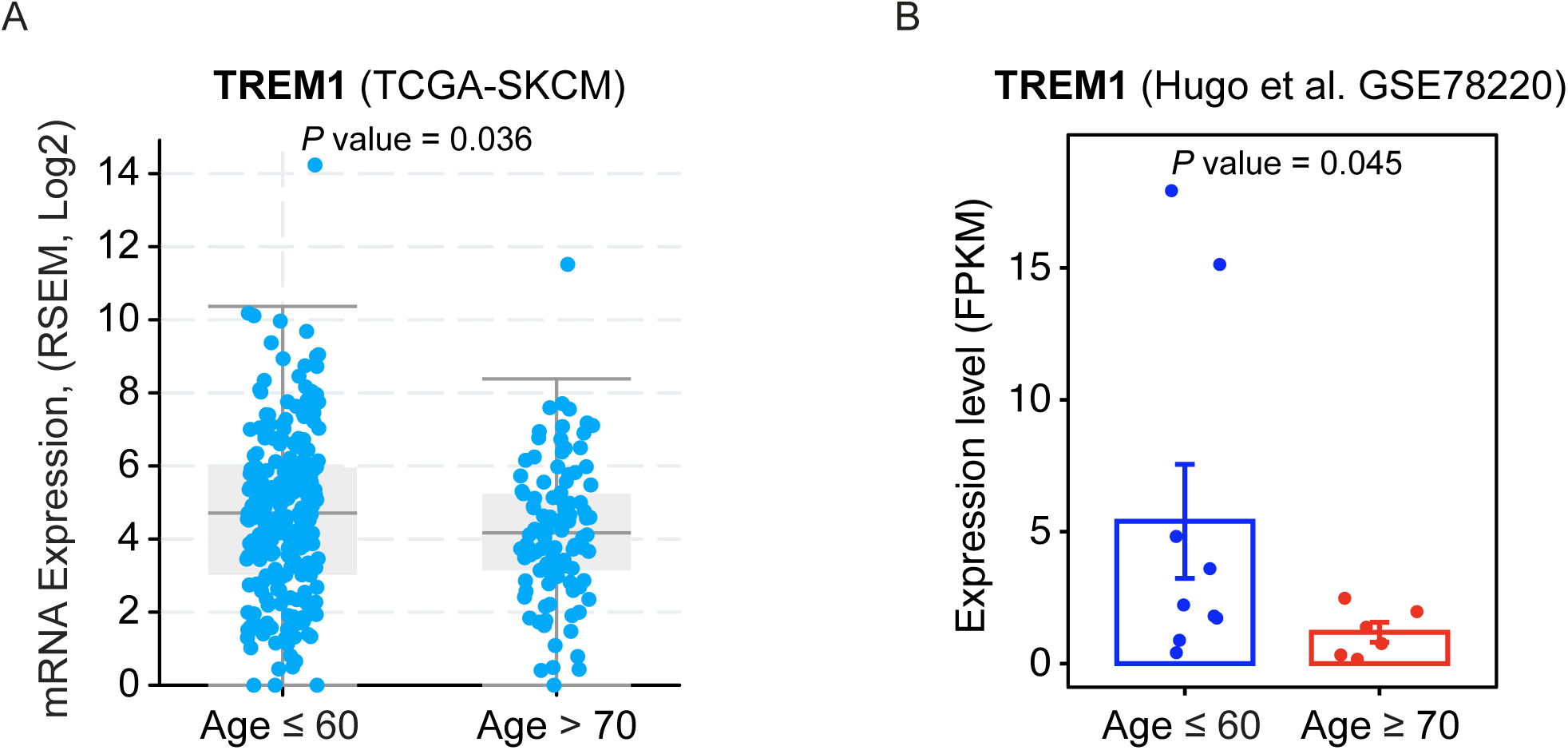
Aging alters TREM1 expression in tumor samples from melanoma patients. (**A**) Expression of TREM1 mRNA in samples of tumors from melanoma patients ≤ 60 and > 70, based on SKCM samples in TCGA dataset. (**B**) Expression of TREM1 mRNA based on the GSE78220 dataset, after stratifying patients by age (≤ 60 and ≥ 70). P-value was computed using Student’s t-test in panel A and one-sided Student’s t-test in panel B.

**Extended Figure 5.**
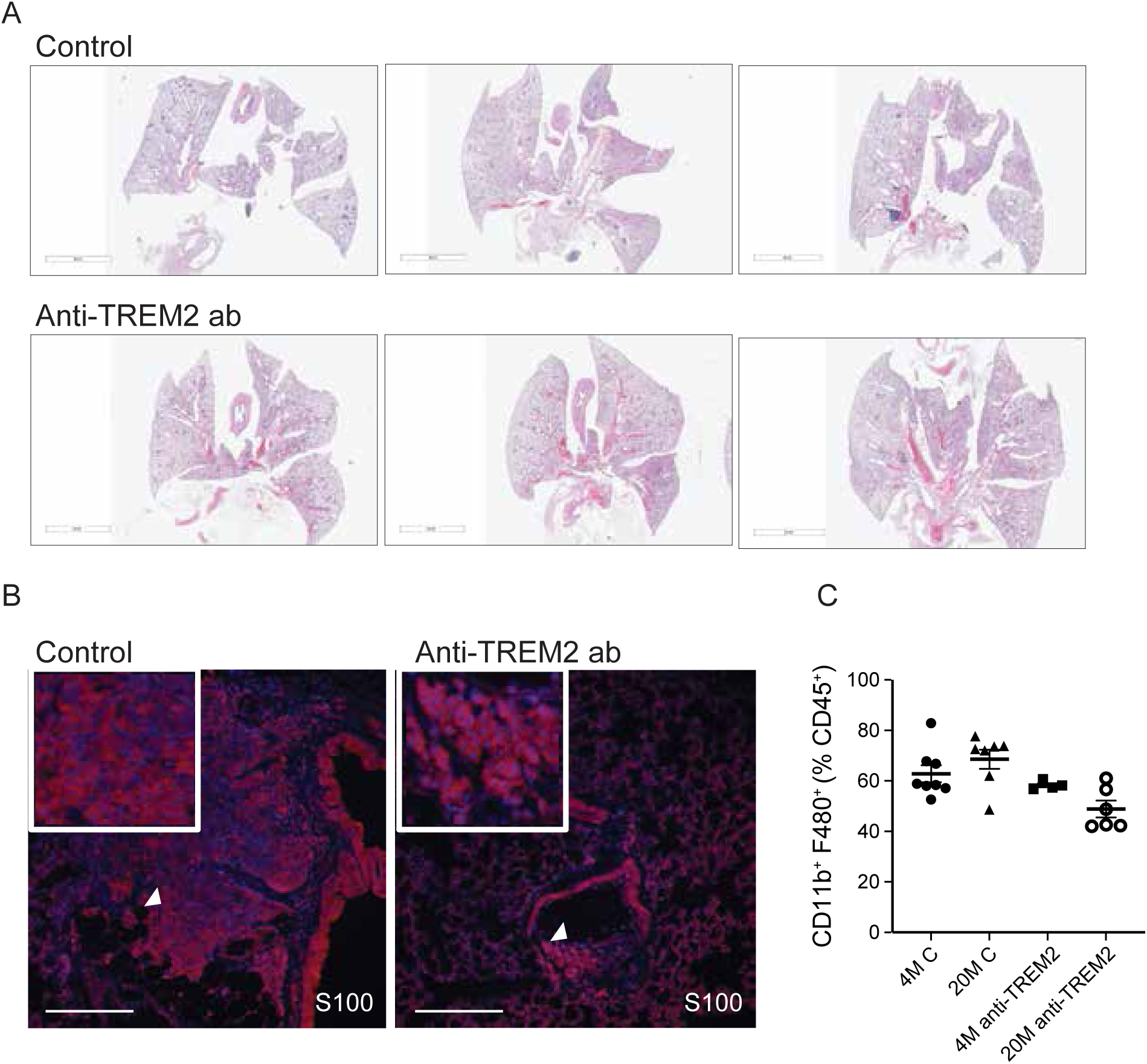
TREM2 inhibitor prevents lung metastasis in aged mice. (**A-B**) YUMM1.7 melanoma cells (500,000) were injected s.c. into the flank of 20M old mice treated either with isotype control or anti-TREM2 antibody and after 22 days lungs were collected and processed to obtain 5 serial sections per lung. (A) Representative H&E staining of lung tissues from indicated. Scale bar, 4 mM. (B) IHC analysis of S100 staining used to detect lung metastasis in samples described above. Scale bar, 100 μM. (**C**) Frequencies of CD11b^+^ and F480^+^ among CD45.2^+^ cells in tumors from 4M and 20M old mice treated with isotype control or anti-TREM2 antibodies. n = 8 for 4M control, n = 4 for 4M TREM2 treated, n = 7 for 20M control, n = 5 for 20M TREM2 treated in panel C. Data in panel C were analyzed by unpaired *t*-test and compared to controls of the same age group.

## Extended data Tables Legends

Extended Table 1. Main UMAP Markers for Figure 1F

Extended Table 2. Macrophages re-clustering markers for Figure 2B

Extended Table 3. DGE of macrophages subclusters 20M versus 4M cluster M1

Extended Table 4. DGE of macrophages subclusters 20M versus 4M cluster M2

Extended Table 5. DGE of macrophages subclusters 20M versus 4M cluster M3

Extended Table 6. DGE of macrophages subclusters 20M versus 4M cluster M4

Extended Table 7. DGE of macrophages subclusters 20M versus 4M cluster M5

Extended Table 8. DGE of macrophages subclusters 20M versus 4M cluster M6

Extended Table 9. DGE of macrophages subclusters 20M versus 4M cluster M7

Extended Table 10. DGE of macrophages subclusters 20M versus 4M cluster M8

Extended Table 11. DGE of macrophages subclusters 20M versus 4M cluster M9

Extended Table 12. CD8 T cells re-clustering markers for Figure 3B

Extended Table 13. Sade-Fedelman Seurat analysis marker genes for Figure 4C

Extended Table 14. Sade-Fedelman Seurat analysis for macrophages marker genes for Figure 5B

Extended Table 15. Genes of Proinflammatory and Tolerogenic signature for Figure 5D

## References

1. Tsai, S., Balch, C. & Lange, J. Epidemiology and treatment of melanoma in elderly patients. Nat Rev Clin Oncol 7, 148–152 (2010).

2. Thompson, J.F. & Williams, G.J. The effect of age on melanoma incidence and prognosis. Aging (Albany NY) 15, 7857–7859 (2023).

3. Fane, M. & Weeraratna, A.T. Normal Aging and Its Role in Cancer Metastasis. Cold Spring Harb Perspect Med 10(2020).

4. Siegel, R.L., Miller, K.D. & Jemal, A. Cancer statistics, 2018. CA Cancer J Clin 68, 7–30 (2018).

5. Hegde, U.P. & Grant-Kels, J.M. Metastatic melanoma in the older patient: special considerations. Clin Dermatol 31, 311–316 (2013).

6. Smetana, K., Jr., et al. Ageing as an Important Risk Factor for Cancer. Anticancer Res 36, 5009–5017 (2016).

7. Lopez-Otin, C., Pietrocola, F., Roiz-Valle, D., Galluzzi, L. & Kroemer, G. Meta-hallmarks of aging and cancer. Cell Metab 35, 12–35 (2023).

8. Franceschi, C., et al. Inflammaging and anti-inflammaging: a systemic perspective on aging and longevity emerged from studies in humans. Mech Ageing Dev 128, 92–105 (2007).

9. Franceschi, C. & Campisi, J. Chronic inflammation (inflammaging) and its potential contribution to age-associated diseases. J Gerontol A Biol Sci Med Sci 69 **Suppl 1**, S4–9 (2014).

10. Fane, M.E., et al. Stromal changes in the aged lung induce an emergence from melanoma dormancy. Nature 606, 396–405 (2022).

11. Bied, M., Ho, W.W., Ginhoux, F. & Bleriot, C. Roles of macrophages in tumor development: a spatiotemporal perspective. Cell Mol Immunol 20, 983–992 (2023).

12. Pan, Y., Yu, Y., Wang, X. & Zhang, T. Corrigendum: Tumor-Associated Macrophages in Tumor Immunity. Front Immunol 12, 775758 (2021).

13. Condeelis, J. & Pollard, J.W. Macrophages: obligate partners for tumor cell migration, invasion, and metastasis. Cell 124, 263–266 (2006).

14. Martin, C.J., Peters, K.N. & Behar, S.M. Macrophages clean up: efferocytosis and microbial control. Curr Opin Microbiol 17, 17–23 (2014).

15. Mills, C.D., Kincaid, K., Alt, J.M., Heilman, M.J. & Hill, A.M. M-1/M-2 macrophages and the Th1/Th2 paradigm. J Immunol 164, 6166–6173 (2000).

16. Ginhoux, F., Schultze, J.L., Murray, P.J., Ochando, J. & Biswas, S.K. New insights into the multidimensional concept of macrophage ontogeny, activation and function. Nat Immunol 17, 34–40 (2016).

17. Cheng, S., et al. A pan-cancer single-cell transcriptional atlas of tumor infiltrating myeloid cells. Cell 184, 792–809 e723 (2021).

18. Mulder, K., et al. Cross-tissue single-cell landscape of human monocytes and macrophages in health and disease. Immunity 54, 1883–1900 e1885 (2021).

19. Peng, J., et al. Single-cell RNA-seq highlights intra-tumoral heterogeneity and malignant progression in pancreatic ductal adenocarcinoma. Cell Res 29, 725–738 (2019).

20. Giladi, A. & Amit, I. Single-Cell Genomics: A Stepping Stone for Future Immunology Discoveries. Cell 172, 14–21 (2018).

21. Moss, C.E., et al. Aging-related defects in macrophage function are driven by MYC and USF1 transcriptional programs. Cell Rep 43, 114073 (2024).

22. Jackaman, C., et al. Aging and cancer: The role of macrophages and neutrophils. Ageing Res Rev 36, 105–116 (2017).

23. Meeth, K., Wang, J.X., Micevic, G., Damsky, W. & Bosenberg, M.W. The YUMM lines: a series of congenic mouse melanoma cell lines with defined genetic alterations. Pigment Cell Melanoma Res 29, 590–597 (2016).

24. Wang, J., et al. UV-induced somatic mutations elicit a functional T cell response in the YUMMER1.7 mouse melanoma model. Pigment Cell Melanoma Res 30, 428–435 (2017).

25. Sudhan, D.R. & Siemann, D.W. Cathepsin L targeting in cancer treatment. Pharmacol Ther 155, 105–116 (2015).

26. Mogilenko, D.A., et al. Comprehensive Profiling of an Aging Immune System Reveals Clonal GZMK(+) CD8(+) T Cells as Conserved Hallmark of Inflammaging. Immunity 54, 99–115 e112 (2021).

27. Sade-Feldman, M., et al. Defining T Cell States Associated with Response to Checkpoint Immunotherapy in Melanoma. Cell 175, 998–1013 e1020 (2018).

28. Hugo, W., et al. Genomic and Transcriptomic Features of Response to Anti-PD-1 Therapy in Metastatic Melanoma. Cell 165, 35–44 (2016).

29. Cheng, X., et al. Systematic Pan-Cancer Analysis Identifies TREM2 as an Immunological and Prognostic Biomarker. Front Immunol 12, 646523 (2021).

30. Homet Moreno, B., et al. Response to Programmed Cell Death-1 Blockade in a Murine Melanoma Syngeneic Model Requires Costimulation, CD4, and CD8 T Cells. Cancer Immunol Res 4, 845–857 (2016).

31. Molgora, M., et al. TREM2 Modulation Remodels the Tumor Myeloid Landscape Enhancing Anti-PD-1 Immunotherapy. Cell 182, 886–900 e817 (2020).

32. Colonna, M. The biology of TREM receptors. Nat Rev Immunol 23, 580–594 (2023).

33. Ford, J.W. & McVicar, D.W. TREM and TREM-like receptors in inflammation and disease. Curr Opin Immunol 21, 38–46 (2009).

34. Tammaro, A., et al. TREM-1 and its potential ligands in non-infectious diseases: from biology to clinical perspectives. Pharmacol Ther 177, 81–95 (2017).

35. Roe, K., Gibot, S. & Verma, S. Triggering receptor expressed on myeloid cells-1 (TREM-1): a new player in antiviral immunity? Front Microbiol 5, 627 (2014).

36. Ajith, A., et al. Targeting TREM1 augments antitumor T cell immunity by inhibiting myeloid-derived suppressor cells and restraining anti-PD-1 resistance. J Clin Invest 133(2023).

37. Sigalov, A.B. A novel ligand-independent peptide inhibitor of TREM-1 suppresses tumor growth in human lung cancer xenografts and prolongs survival of mice with lipopolysaccharide-induced septic shock. Int Immunopharmacol 21, 208–219 (2014).

38. Li, R.Y., et al. TREM2 in the pathogenesis of AD: a lipid metabolism regulator and potential metabolic therapeutic target. Mol Neurodegener 17, 40 (2022).

39. Guo, X., et al. TREM2 promotes cholesterol uptake and foam cell formation in atherosclerosis. Cell Mol Life Sci 80, 137 (2023).

40. Wolf, E.M., Fingleton, B. & Hasty, A.H. The therapeutic potential of TREM2 in cancer. Front Oncol 12, 984193 (2022).

41. Katzenelenbogen, Y., et al. Coupled scRNA-Seq and Intracellular Protein Activity Reveal an Immunosuppressive Role of TREM2 in Cancer. Cell 182, 872–885 e819 (2020).

42. Nalio Ramos, R., et al. Tissue-resident FOLR2(+) macrophages associate with CD8(+) T cell infiltration in human breast cancer. Cell 185, 1189–1207 e1125 (2022).

43. Tan, J., et al. TREM2(+) macrophages suppress CD8(+) T-cell infiltration after transarterial chemoembolisation in hepatocellular carcinoma. J Hepatol 79, 126–140 (2023).

44. Lei, X., Gou, Y.N., Hao, J.Y. & Huang, X.J. Mechanisms of TREM2 mediated immunosuppression and regulation of cancer progression. Front Oncol 14, 1375729 (2024).

45. Binnewies, M., et al. Targeting TREM2 on tumor-associated macrophages enhances immunotherapy. Cell Rep 37, 109844 (2021).

46. Park, M.D., et al. TREM2 macrophages drive NK cell paucity and dysfunction in lung cancer. Nat Immunol 24, 792–801 (2023).

47. Xiong, D., Wang, Y. & You, M. A gene expression signature of TREM2(hi) macrophages and gammadelta T cells predicts immunotherapy response. Nat Commun 11, 5084 (2020).

48. Chen, A.C.Y., et al. The aged tumor microenvironment limits T cell control of cancer. Nat Immunol 25, 1033–1045 (2024).

49. Han, S., Georgiev, P., Ringel, A.E., Sharpe, A.H. & Haigis, M.C. Age-associated remodeling of T cell immunity and metabolism. Cell Metab 35, 36–55 (2023).

50. Drijvers, J.M., Sharpe, A.H. & Haigis, M.C. The effects of age and systemic metabolism on anti-tumor T cell responses. Elife 9(2020).

51. Bovenschen, N., et al. Granzyme K displays highly restricted substrate specificity that only partially overlaps with granzyme A. J Biol Chem 284, 3504–3512 (2009).

52. Tiberti, S., et al. GZMK(high) CD8(+) T effector memory cells are associated with CD15(high) neutrophil abundance in non-metastatic colorectal tumors and predict poor clinical outcome. Nat Commun 13, 6752 (2022).

53. Dorshkind, K., Montecino-Rodriguez, E. & Signer, R.A. The ageing immune system: is it ever too old to become young again? Nat Rev Immunol 9, 57–62 (2009).

54. Sidler, C., et al. Immunosenescence is associated with altered gene expression and epigenetic regulation in primary and secondary immune organs. Front Genet 4, 211 (2013).

55. Szarc vel Szic, K., Declerck, K., Vidakovic, M. & Vanden Berghe, W. From inflammaging to healthy aging by dietary lifestyle choices: is epigenetics the key to personalized nutrition? Clin Epigenetics 7, 33 (2015).

56. Franceschi, C., Garagnani, P., Vitale, G., Capri, M. & Salvioli, S. Inflammaging and ’Garb-aging’. Trends Endocrinol Metab 28, 199–212 (2017).

57. Sambrano, G.R. & Steinberg, D. Recognition of oxidatively damaged and apoptotic cells by an oxidized low density lipoprotein receptor on mouse peritoneal macrophages: role of membrane phosphatidylserine. Proc Natl Acad Sci U S A 92, 1396–1400 (1995).

58. Wang, Y., et al. TREM2 lipid sensing sustains the microglial response in an Alzheimer’s disease model. Cell 160, 1061–1071 (2015).

59. Atagi, Y., et al. Apolipoprotein E Is a Ligand for Triggering Receptor Expressed on Myeloid Cells 2 (TREM2). J Biol Chem 290, 26043–26050 (2015).

60. Bailey, C.C., DeVaux, L.B. & Farzan, M. The Triggering Receptor Expressed on Myeloid Cells 2 Binds Apolipoprotein E. J Biol Chem 290, 26033–26042 (2015).

61. Cochran, A.J. & Wen, D.R. S-100 protein as a marker for melanocytic and other tumours. Pathology 17, 340–345 (1985).

62. Dobin, A., et al. STAR: ultrafast universal RNA-seq aligner. Bioinformatics 29, 15–21 (2013).

63. Frankish, A., et al. GENCODE: reference annotation for the human and mouse genomes in 2023. Nucleic Acids Res 51, D942–D949 (2023).

64. Hao, Y., et al. Integrated analysis of multimodal single-cell data. Cell 184, 3573–3587 e3529 (2021).

65. Butler, A., Hoffman, P., Smibert, P., Papalexi, E. & Satija, R. Integrating single-cell transcriptomic data across different conditions, technologies, and species. Nat Biotechnol 36, 411–420 (2018).

66. Hoadley, K.A., et al. Cell-of-Origin Patterns Dominate the Molecular Classification of 10,000 Tumors from 33 Types of Cancer. Cell 173, 291–304 e296 (2018).

67. Cerami, E., et al. The cBio cancer genomics portal: an open platform for exploring multidimensional cancer genomics data. Cancer Discov 2, 401–404 (2012).

68. Wickham H. Data analysis. In: ggplot2: elegant graphics for data analysis. Springer; 2016. p. 189–201. doi: doi.org/10.1007/978-0-387-98141-3.

